# Intervention reducing malaria parasite load in vector mosquitoes: no impact on *Plasmodium falciparum* extrinsic incubation period and the survival of *Anopheles gambiae*

**DOI:** 10.1101/2022.12.28.522076

**Authors:** Edwige Guissou, Dari Frédéric Da, Domombabele François de Sales Hien, Koudraogo Bienvenu Yameogo, Serge Rakiswende Yerbanga, Anicet Georges Ouédraogo, Kounbobr Roch Dabiré, Thierry Lefèvre, Anna Cohuet

## Abstract

In the fight against malaria, transmission blocking interventions (TBIs) are promising approaches to complement conventional tools. They aim to prevent the infection of the vectors and thereby reduce the subsequent exposure of a human population to infectious mosquitoes. The effectiveness of these approaches has been shown to depend on the initial intensity of infection in mosquitoes, often measured as the mean number of oocysts resulting from an infectious blood meal prior to intervention. In mosquitoes exposed to a high infection load, current TBI candidates are expected to be ineffective at completely blocking infection but will decrease parasite load and therefore, potentially also affect key parameters of vector transmission. The present study aims to investigate the consequences of changes in oocyst intensity on downstream parasite development and mosquito survival. To address this, we experimentally produced different infection intensities in *Anopheles gambiae* females by diluting gametocytes from natural *Plasmodium falciparum* isolates and used a newly developed non-destructive method based on the exploitation of mosquito sugar feeding to track parasite and mosquito life history traits throughout sporogonic development. Our results indicate that *P. falciparum* extrinsic incubation period (EIP) and mosquito survival did not vary with parasite density but differed significantly between parasite isolates. Our results do not identify here unintended consequences of the decrease of parasite loads in mosquitoes on the parasite incubation period or on mosquito survival, two key parameters of vectorial capacity, and hence support the transmission blocking strategies to control malaria.

**Author summary:** In the fight against malaria, it is recognized that the use of several complementary strategies is necessary to significantly reduce transmission and improve human health. Among these, transmission blocking strategies aim at blocking the development of the parasites within the mosquito vectors. This approach should prevent infection in most of mosquitoes feeding on infectious hosts and thus block the transmission. However, in some cases it may only reduce parasite load without fully clearing the infection. Here we identified potential risks: if reducing parasite load would reduce the incubation period of the parasite in the mosquitoes or increase the longevity of the mosquitoes, undesirable consequences may occur with an increased efficiency of these mosquitoes to transmit parasites to human. We therefore tested these hypotheses and experimentally produced different infection loads in vector mosquitoes *Anopheles gambiae* by using dilutions of *Plasmodium falciparum* isolates from naturally infected human donors. We observed that the longevity of the mosquitoes and the incubation period of the parasites were not affected by the parasite load. This is not consistent with the unintended risks that we investigated and thus strengthens the potential of transmission blocking interventions in the toolbox to combat malaria.

## Introduction

Despite significant progress in the fight against malaria in the last two decades, nearly half of the world’s population remain at risk of contracting the disease. The African region is the most affected, accounting for 94% of the global malaria burden [1]. Malaria control mainly relies on the use of antimalarial drugs, with a great contribution of artemisinin-based combination therapies, and vector control with the use of long-lasting insecticidal nets and indoor residual sprayings. These tools allowed the significant reduction of malaria incidence and mortality that occurred since the beginning of the century, but this decline has worryingly stalled in some countries and even reversed in some others during recent years with the spread of drug-resistance among parasites [2,3] and insecticide resistance in the main mosquito vectors [4,5].

As a complement to conventional tools that target the parasite in humans or seek to kill mosquito vectors, targeting parasite within the mosquito appears to be a promising approach [6–8]. These approaches, known as transmission blocking interventions (TBIs), can use drugs [9,10] or vaccines [11–14] to impede the infection of the vectors and thereby reduce the subsequent exposure of a human population to infectious mosquitoes. The efficiency of such approaches has been shown to depend on the initial intensity of infection in the mosquito, often measured as the mean number of oocysts resulting from an infectious blood meal before the intervention [15–17]. Indeed, when the intensity of infection is moderate or low (< 5 oocysts per mosquito) as often found in naturally infected mosquitoes [18–20], transmission-blocking strategies will be much more effective in reducing the prevalence of infection in mosquitoes compared to situations where mosquitoes carry higher parasite intensities. However, in nature, it has been shown that the distribution of oocysts is highly overdispersed, with a significant proportion of infected mosquitoes carrying dozens of oocysts and few mosquitoes harboring very high oocyst densities (>50 oocysts per mosquito) [21]. In mosquitoes carrying high infection load, it is then expected that imperfect TBIs will reduce the number of oocysts but will be ineffective at completely blocking infection.

In studies that evaluate the efficiency of TBIs, the effect of the intervention on infection intensity, or transmission reducing activity (TRA), is used as a predictor of the transmission blocking activity (TBA), the efficacy at blocking the infection [16] but the consequences of the decrease of infection load among mosquitoes that remain infected despite the intervention is barely considered. One reason is that malaria epidemiological studies typically use the proportion of mosquitoes carrying sporozoites (the sporozoite rate among wild mosquito vectors) as an indicator of exposure risk to the human population and implicitly assume that these mosquitoes are equally infectious irrespective of the parasite load they carry (e.g. [22]). However, thinner considerations of epidemiological parameters suggest that, reducing the intensity of infection in mosquitoes, without successfully clearing the infection, may have indirect consequences on transmission. First, some studies showed that mosquitoes heavily infected with sporozoites are more likely to induce successful subsequent infections than counterparts infected with lower sporozoite loads [23–26]. Considering a direct correlation between the oocyst intensity when they all have ruptured and sporozoite intensity in salivary glands [27,28], this suggests that TBIs may be beneficial beyond the effect on infection prevalence and that a reduction of infection intensity in mosquitoes, even without clearing infection, may also decrease mosquito to human transmission. Second, several evolutionary considerations and experimental observations suggest that density of parasites may affect their dynamics of development [29,30] and highlight other possible consequences of imperfect TBIs on transmission. On the one hand, the decrease in intensity could be accompanied by a decrease in the multiplicity of infection, and thus a release of competition between parasite genotypes [31,32]. The parasites might then decrease their investment in the production of transmissible stages and thus lengthen the time between the infectious blood meal and when the mosquito becomes infectious to the vertebrate, i.e. the extrinsic incubation period (EIP). In the same way, stochastic models show that a reduced oocyst number decreases the chance that one has already released some sporozoites and induces a longer EIP [33]. On the other hand, the limited resources in the mosquito may become less restrictive when infection intensity decreases which may accelerate the development of parasites from oocysts to sporozoites and decrease the EIP [34,35]. Because the EIP in malaria parasites is almost as long as life expectancy of malaria vectors, this parameter is a key factor of transmission [36] and the potential consequences of a malaria TBIs on EIP remains to be investigated. Closely related to EIP, mosquito survival is also a parameter of vectorial capacity, which drastically influences the intensity of transmission and deserves attention. The effect of *Plasmodium* infection on survival of its natural vectors appeared to be dependent on environmental conditions [37–40]. Regarding the intensity of the infection, some studies have shown that oocysts intensity negatively affects mosquitoes survival [41,42]. Lower *Plasmodium* infection intensity in mosquitoes, in consequences to TBIs, may then decrease virulence of parasites within its vector and in that way facilitate its transmission. It is therefore important to understand the effect of parasite density on this key transmission parameter and investigate whether TBIs would unintentionally increase the life expectancy of the vectors that remain infected by reducing the infection load and thus possibly facilitate successful and sustained transmission of the pathogen.

Because imperfect TBIs will affect parasite intensity in mosquitoes and possibly affect key parameters of vectorial transmission, the present study aims at investigating the consequences of changes in oocyst intensity on downstream parasite development and mosquito survival. We experimentally produced different intensities of infections in *Anopheles gambiae* females by diluting gametocytes from natural isolates of *Plasmodium falciparum* and used a newly developed non-destructive method [43] based on the exploitation of mosquito sugar feeding to track parasite and mosquito life history traits throughout sporogonic development.

## Results

### Effect of infectious blood dilution on the prevalence and intensity of infection in mosquitoes

#### • Infection prevalence and intensity in mosquito gut at 7 day post blood meal (dpbm)

*An. gambiae* females were experimentally infected with the blood from one of three naturally infected gametocytes carriers (parasite isolates A, B and C) in Burkina Faso. The gametocyte-infected blood of each carrier was diluted to experimentally reduce the density of infectious gametocytes and create a range of parasite loads in mosquitoes. The midguts of 193 females were dissected on 7 dpbm for oocyst observation, among which 132 were positive to *P. falciparum* (68.4%). Dilution had no significant effect on the infection prevalence (LRT X^2^_1_=1.8, P=0.176, Figure 1a) but had the intended effect with a strong reduction in infection intensity among oocyst-infected mosquitoes throughout the dilution range (LRT X^2^_1_=7.1, P=0.008, Fig 1b).

**Fig 1:**
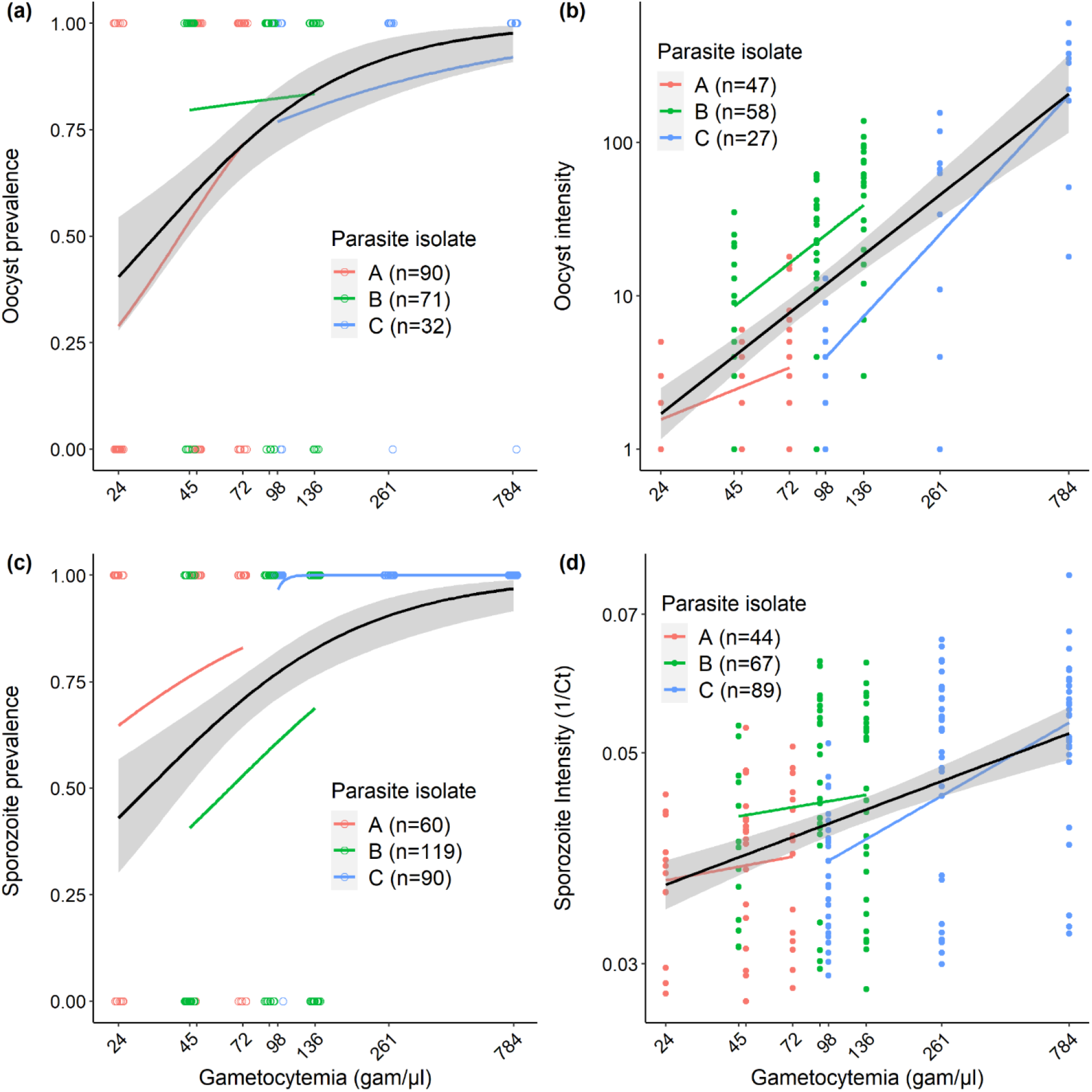
Effect of infectious blood dilution on the prevalence and intensity of infection in mosquitoes. (a) Oocyst prevalence 7 dpbm (number of mosquitoes with at least one oocyst in their midguts out of the total number of mosquitoes dissected) as a function of gametocytemia (the estimated number of infectious gametocytes per microliter of blood in each of the dilution group: 24, 45, 48, 72, 91, 98, 136, 261 and 784 gametocytes/μl of blood, note that to avoid overlapping of the x-axis labels, the concentrations of 48 and 91 gam/μl are not indicated). Each colored circle represents a dissected mosquito (red color: parasite isolate A with an initial gametocytemia of 72, green color: parasite isolate B with initial gametocytemia of 136, and blue color: parasite isolate C with an initial gametocytemia of 784). The colored lines represent the best-fit logistic growth curves for each parasite isolate, while the black line (± se) for all data regardless of isolate origin. Note that the x-axis is on a log10 scale. (b) Oocyst intensity 7 dpbm (number of oocysts in infected mosquitoes) as a function of gametocytemia. Each colored circle represents a *P. falciparum* oocyst-positive midgut. The colored lines represent the linear relationship for each parasite isolate, while the black line (± se) for all data regardless of isolate origin. Note that the x- and y-axes are on a log10 scale. (c) Sporozoite prevalence (number of mosquitoes with heads/thoraces detected positive to *P. falciparum* at death of the individual using qPCR out of the total number of tested heads/thoraces) as a function of gametocytemia. Each colored circle represents a tested head/thorax. The x-axis is on a log10 scale. (d) Amount of parasite DNA in mosquito heads/thoraces expressed as the inverse of the qPCR cycle threshold (1/Ct, the higher the inverse of threshold cycle, the higher the intensity of infection). For each mosquito, 1/Ct value is the average over 4 to 6 technical replicates. The x- and y-axes are on a log10 scale. Each colored circle represents a *P. falciparum* positive head/thorax using qPCR.

#### • Infection prevalence and intensity in mosquito head/thoraces upon death

At 7 dpbm, 269 *An. gambiae* females that were challenged with either parasite isolate A, B or C and that had received the different dilution treatments (1/1, 1/3, 2/3 for parasite isolates A and B and 1/1, 1/3, 1/8 for parasite isolate C), were individually placed in tubes for saliva collection on cotton balls soaked in a 10% glucose solution. Upon mosquito death, the amount of parasite DNA in the heads/thoraxes of the females used to collect saliva was assessed using qPCR. Of a total of 269 *An. gambiae* females placed in individual tubes, 201 (75%) were found positive to *P. falciparum* by qPCR on the heads/thoraces at death of the mosquito. The proportion of sporozoite-infected mosquitoes significantly increased with the density of infectious gametocytes (LRT X^2^_1_=4.7, P=0.030, Fig 1c). The global percentage of positive heads/thoraces reaches 88% (151out of 172) when females that died before 14 dpbm (the time generally considered for sporozoites to have invaded mosquito salivary glands) were excluded. Consistent with observations made in mosquito’s midguts, the amount of parasite DNA among infected mosquito’s heads and thoraces showed a positive relationship with gametocytemia (LRT X^2^_1_=24.3, P< 0.001, Fig 1d).

### Effect of parasite density on EIP

The presence of parasite DNA in the cotton balls used to collect saliva from infected mosquitoes (n=201) was examined. A total of 1 997 cotton balls were analyzed and individual EIP was defined as the time between the infectious blood meal and the first day of positive qPCR detection of *P. falciparum* from a cotton wool substrate for a given infected female. Of the 201 females with infected head/thorax, 102 (50.7%) generated at least one cotton ball containing detectable traces of parasite DNA. The infected females that did not produce any positive cottons over their lifespan were excluded from the analysis because no EIP values can be derived from these samples. The first positive cottons occurred on 9 dpbm in all dilution treatment.

There was no effect of gametocytemia on EIP (*LRT X^2^_1_* = 0.9, P = 0.341, Fig 2a). Similar results were obtained when the explanatory variable, the gametocytemia, was substituted by the actual individual infection intensity found in the head thorax of each female upon death (*LRT X^2^]* = 1.0, P = 0.317, Fig 2b). However, there was a significant effect of parasite isolate (*LRT X^2^_2_* = 32.8, P < 0.001) with an EIP_50_ of 16, 14 and 12 days for isolate A, B and C, respectively (Fig 2c).

**Fig 2:**
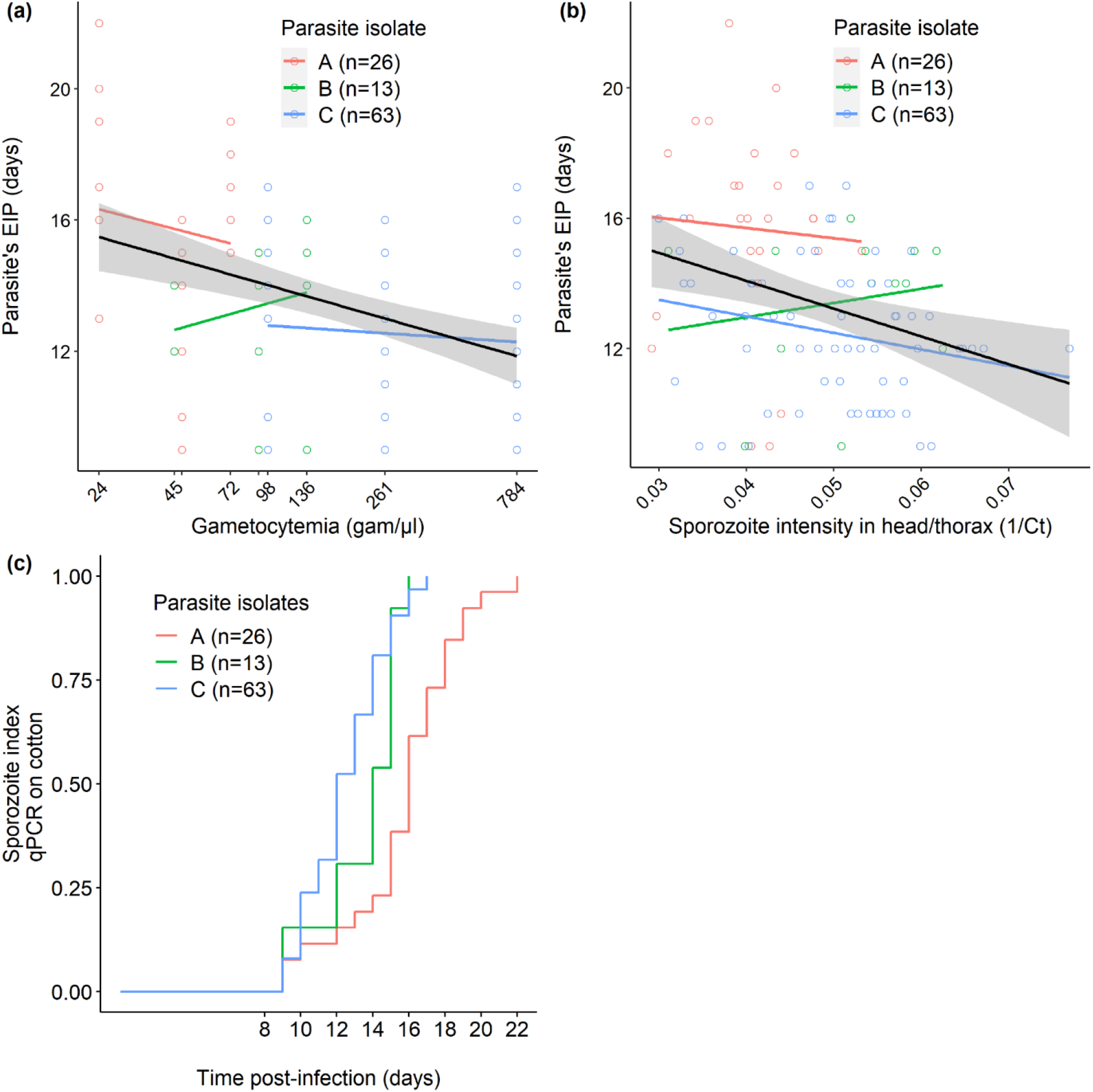
Relationship between parasite density and EIP. (a) EIP (the time between the infectious blood meal and the detection of *P. falciparum* in mosquito saliva collected from cotton balls) as a function of gametocytemia (the number of gametocytes per microliter of blood in each of the dilution group: 24, 45, 48, 72, 91, 98, 136, 261 and 784 gametocytes/μl of blood, note that to avoid overlapping of the x-axis labels, the concentrations of 48 and 91 gam/μl are not indicated). Each colored circle represents an infected mosquito from which EIP was measured (red color: parasite isolate A with an initial gametocytemia of 72, green color: parasite isolate B with initial gametocytemia of 136, and blue color: parasite isolate C with an initial gametocytemia of 784). The x-axis is on a log10 scale. (b) EIP as a function of 1/Ct in mosquito infected heads/thoraces extracts (the higher the 1/Ct value, the higher the infection intensity). For each cotton ball, 1/Ct value is the average over 3 technical replicates. The colored lines in panels’ a and b represent the linear relationship for each parasite isolate, while the black line (± se) for all data regardless of isolate origin. (c) Kaplan–Meier curves representing the temporal dynamics of sporozoite detection in cotton balls used to collect saliva from individual mosquitoes fed on each parasite isolate.

### Effect of parasite density on mosquito survival

The survival was monitored daily for the 269 females placed in individual tubes, including 201 infected with *P. falciparum* and 68 fed on infectious blood but that did not develop an infection. Infected females showed a better survival in individual tubes compared to the ones that did not become infected (*LRT X^2^_1_* = 14.2, P < 0.001, Fig 3a). No interaction between infection status and gametocytemia on mosquito survival was found (*LRT X^2^_1_* = 0, P = 0.968, Fig 3b). There was no effect of gametocytemia on the lifespan of infected mosquito females (*LRT X^2^_1_* = 0.1, P =0.783, Fig 3c) and no effect of infection intensity in mosquito’s head/thorax on the lifespan of infected mosquito females (*LRT X^2^_1_* = 1.9, P =0.165, Fig 3d). Finally, there was a strong lifespan variation among infected females depending on parasite isolates (*LRT X^2^_1_* = 46.6, P <0.001, Fig 3e), with a median longevity of 25, 15 and 18 days in infected mosquitoes fed respectively on isolate A, B and C.

**Fig 3:**
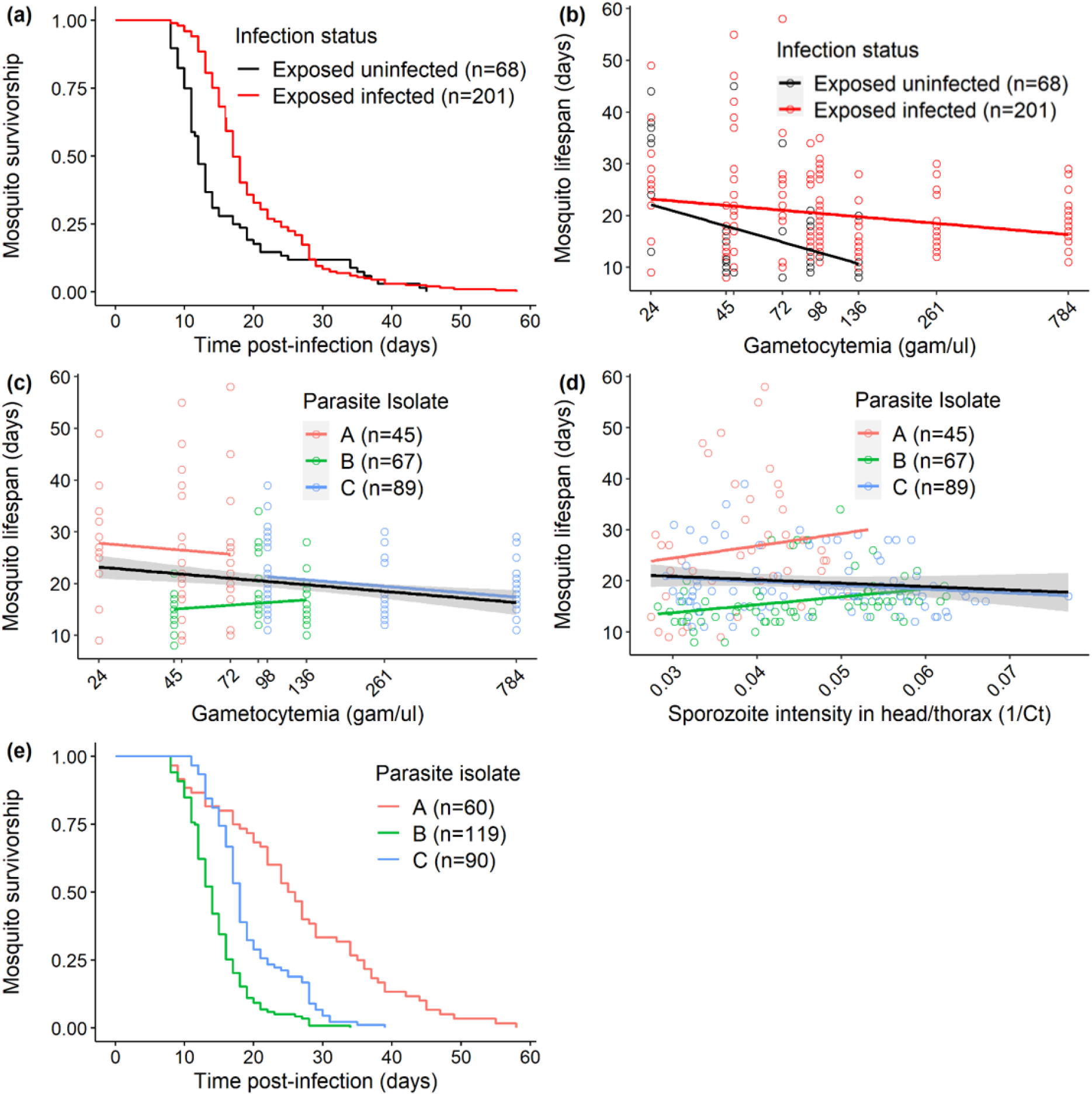
Relationship between parasite density and mosquito longevity. (a) Kaplan–Meier curves representing the survival in days post blood meal for each infection status (red color: mosquitoes exposed to infectious blood and having developed *Plasmodium*, black color: mosquitoes exposed to infectious blood and which remained uninfected). (b) Mosquito longevity in days as a function of gametocytemia (the number of gametocytes per microliter of blood in each of the dilution group: 24, 45, 48, 72, 91, 98, 136, 261 and 784 gametocytes/μl of blood, note that to avoid overlapping of the x-axis label, the concentrations of 48 and 91 gam/μl are not indicated). Each colored circle represents a mosquito exposed to the infectious blood (red color: infected mosquitoes, black color: mosquitoes that remained uninfected). The x-axis is on a log10 scale. (c) Mosquito longevity in days as a function of gametocytemia (the number of gametocytes per microliter of blood in each of the dilution group: 24, 45, 48, 72, 91, 98, 136, 261 and 784 gametocytes/μl of blood, note that to avoid overlapping of the x-axis label, the concentrations of 48 and 91 gam/μl are not indicated). Each colored circle represents a mosquito exposed to one of the three parasite isolates (red color: parasite isolate A with an initial gametocytemia of 72, green color: parasite isolate B with initial gametocytemia of 136, and blue color: parasite isolate C with an initial gametocytemia of 784). The x-axis is on a log10 scale. (d) mosquito longevity in days as a function of 1/Ct in mosquito infected heads/thoraces extracts (the higher the 1/Ct value, the higher the infection intensity). For each cotton ball, 1/Ct value is the average over 3 technical replicates in two qPCR machines. Each colored circle represents an infected mosquito. The colored lines in panels’ b and c represent the linear relationship for each parasite isolate, while the black line (± se) for all data regardless of isolate origin. (e) Kaplan–Meier curves representing the survival in days post blood meal for each parasite isolate.

## Discussion

In the present study, we questioned the importance of infection load in malaria-infected mosquitoes. We investigated the relationships between *P. falciparum* gametocyte densities in infectious blood and subsequent transmission parameters in mosquitoes, including infection prevalence, infection intensity at oocyst and sporozoite stages, and more originally the time it takes for the mosquitoes to become infectious (parasite’s EIP) and their survival. In particular, EIP and mosquito survival are key parameters of transmission [44–46] and we explored the extent to which any intervention affecting the intensity of infection could affect them. Because infection outcome in mosquitoes depends on various parameters such as uncertainty in the gametocytes quantification [47–51], gametocytes maturity and sex ratio [49,52–54], genetics [55–58], parasite multiplicity of infection [54,59], as well as environmental conditions [39,60] we generated experimental ranges of parasite loads from infectious blood samples so that for a given parasite isolate, only the infectious gametocyte density varied. To do this, we diluted field collected gametocyte containing blood isolates by an own fraction of the blood sample that was exposed to heat inactivation and we used the range of dilutions to expose mosquitoes in controlled conditions by DMFAs.

The range of *P. falciparum* infectious gametocyte densities generated by dilution resulted in proportional oocyst intensities of infection in *An. gambiae* mosquitoes. This relationship between gametocyte density and oocyst load in mosquitoes was previously demonstrated in study using the same dilution protocol [61]. Here we observed similar relationships between gametocyte densities and sporozoite load in mosquito head/thorax extracts, consistent with linear relation between oocyst number and sporozoite load in salivary glands [27,28]. Surprisingly, the sporozoite prevalence we observed by qPCR in head/thorax extracts of mosquitoes from 14 dpbm (88%) were higher than oocyst prevalence in midguts at 7 dpbm (68.4%), suggesting that qPCR for sporozoite detection could be more sensitive than microscope detection of oocysts, may produce false positives or that infected females could survive better than exposed but non-infected females. These three hypotheses are not mutually exclusive; the fact that we observed an important proportion of females positive for *P. falciparum* in the heads and thoraces that never produced a positive cotton (99 individuals out of 201) suggests that the inclusion of 4 to 6 technical replicates may have overestimated the proportion of positive carcasses by favoring false positives and support the second hypothesis. Moreover, the survival recording revealed that females exposed to infectious blood but that did not develop an infection lived shorter than their infected counterparts (Fig 3a), therefore providing support for the third hypothesis. As our study focused on life history traits of mosquitoes from which *P. falciparum* were detected in expelled saliva, the discrepancies between the ratio of head and thorax positive mosquitoes compared to oocyst positive and cotton positive detection did not affect our conclusions.

Until recently, studying life history traits of infected mosquitoes and in particular following the dynamic of parasites in mosquitoes required to dissect a large number of individuals and did not allow recording all variables from a given female. Here, we took advantage of the recent development of a non-destructive technique to measure the presence of sporozoites in the saliva of mosquitoes as a proxy of their infectivity without sacrificing them so that the different parameters could be measured in the same individuals, allowing to highlight trade-offs [43]. Our results show that the manipulation of gametocyte density and subsequently of oocyst and sporozoite densities has no effect on the parasite’s extrinsic incubation period. Although not significant, mosquitoes exposed to the highest gametocyte densities and carrying the highest sporozoite loads had slightly shorter EIP. This could be due to a previously identified bias in our non-destructive method of sporozoite detection. Indeed, this technique, based on the detection of sporozoite DNA in absorbent cotton on which females have come to take a sugar meal and have left saliva infected with *P falciparum*, is subject to limitations related to detection thresholds. A consequence is that higher sporozoite loads are more likely to be detected so that our technique may have overestimated the extrinsic incubation time in mosquitoes carrying low sporozoite loads [43]. Thus, despite the observed trend, which remains non-significant and likely to be due to the expected technical bias, our results highlight that the EIP does not depend on the parasite density in the system tested here. These are not consistent with the predictions that lower infection intensities in mosquitoes would limit the competition for resources between parasites and therefore speed up their development (i.e. shorter EIP). Recent results obtained by using an inbred parasite strain [34] in contrast supported this hypothesis but our results suggest more complex relationships where, for instance, different parasite genotypes or multiplicity of infection could induce variable effects on EIP.

Regarding the survival of infected mosquitoes, one expectation was that higher parasite loads could affect the longevity of females compared to females infected with lower loads [41,42,62]. If the question of the effect of *Plasmodium* infection on mosquito survival is under debate for decades, the effect of dose-dependent effect remained to be explored. Our observations did not show such a correlation with mosquito lifespan not dependent on parasite load. Besides, it appears that the females exposed to the infection but which remained uninfected displayed reduced longevity compared to infected ones. This suggests either a mutualist interaction between *P. falciparum* and *An. gambiae*, with a benefice of being infected for the host, or a cost of resistance that would reduce the fitness of resistant hosts associated to a tolerance of the infected ones. Our present data do not allow to discriminate between the two hypotheses as a non-exposed mosquito control group would have been needed to determine if infection increases/maintains the host’s lifespan or resistance a reduced one. In addition, trade-offs between lifespan and fecundity are expected [63] and further studies based on a more complete picture of individual mosquitoes’ fitness are still needed. However, to date several previous studies suggest that resisting the infection induced a reduced survival of mosquitoes, consistent with a cost of resistance, although high variance was found between assays [40,64,65]. Therefore, to better depict the interactions between malaria vectors and parasites, future studies, using the non-destructive detection of parasite at the mosquito individual level, will still be needed to examine possible associations between vector survival and fecundity and parasite load and EIP.

Regardless of the intensity of the parasites, our study reveals that EIP and survival greatly varied depending on the parasite isolates and assay replicates, which are confounded here. In regions with high malaria transmission such as Burkina Faso, previous studies have shown a high genetic diversity in *P. falciparum* isolates [66,67]. Our results therefore suggest that there is a variation in the EIP and mosquito survival depending on the genetics of the parasite isolates. It can be hypothesized that isolates with a multiplicity of infection would favor competition between genotypes and possibly the rapidity of sporogony and virulence [29,32,68]. Studies are therefore ongoing to determine the effect of the genetic diversity of parasite isolates on their EIP and their effect on mosquito survival.

Our study provides findings on the effects of *Plasmodium* parasite load in mosquito vectors on the life history traits of the mosquito and the parasite that could influence transmission. In this sense, it sheds light on the potential consequences of interventions against malaria, which, if they do not always succeed in completely blocking the transmission of the parasite, could result in a decrease of the parasite load in mosquitoes. If a decrease in parasite load in the mosquito resulted in a strong decrease in EIP for the parasite or increase of longevity for the vectors, the consequences in terms of transmission could be counterproductive, with an increase in the risk of exposure to infecting bites. Our results do not show such consequences and therefore do not identify a risk associated to the decrease of the parasite load in the mosquito on the parasite extrinsic incubation time in the mosquito or their survival, and support here the strategies of blocking the transmission of malaria.

## Materials and Methods

### Mosquitoes

In this study, we used an outbred colony of *An. gambiae* that was established in 2008 and repeatedly replenished with F1 from wild-caught mosquito females collected in Soumousso, (11°23’14”N’ 4°24’42”W), 40 km from Bobo Dioulasso, south-western Burkina Faso (West Africa). To do so, field collected fed or gravid *Anopheles* females, morphologically identified as belonging to the *An. gambiae* complex, were further identified by using a SINE-PCR [69] before pooling the eggs of *An. gambiae s.s.*. Mosquitoes were then held in 30 × 30 × 30 cm mesh-covered cages and maintained under standard insectary conditions (27 ± 2 °C, 70 ± 5% HR, 12:12 LD) in the IRSS (Institut de Recherche en Sciences de la Santé) laboratory in Bobo Dioulasso.

### *P. falciparum* natural isolates, infectious gametocytes dilution and mosquito infection

*Anopheles gambiae* female mosquitoes were exposed to blood samples from donors naturally infected with *P. falciparum* gametocytes using direct membrane feeding assay (DMFA) as described previously [70] and with a dilution procedure [61].

Briefly, thick blood smears were carried out from volunteers among 5-12 year-old schoolchildren in villages around Bobo-Dioulasso, air-dried, Giemsa-stained, and examined microscopically for the presence of *P. falciparum*. Asexual trophozoites parasite stages were counted against 200 leukocytes, while mature gametocyte stages were counted against 1,000 leukocytes and parasite densities were estimated on the basis of an average of 8,000 leukocytes/ml. Children with an asexual parasitaemia of > 1,000 parasites per microliter were treated according to national guidelines. Blood samples of three asymptomatic *P. falciparum* gametocyte carriers (called isolates A, B and C) were collected by venipuncture in heparinized tubes and their plasma was replaced by AB serum from a European donor. These blood samples underwent dilution series. The dilution consisted in heating at 45°C for 20 minutes part of each blood sample to inactivate the infectivity of gametocytes [71] and using this non-infectious blood to reduce the density of infectious parasites for each isolate. The blood isolate A, with 72 gametocytes/μl of blood, was treated to obtain three dilution factors, namely 1/1 (undiluted blood, 72 gametocytes/μl), 2/3 (48 gametocytes/μl) and 1/3 (24 gametocytes/μl). Isolate B with 136 gametocytes/μl of blood was diluted according to the same dilution factors than isolate A: 1/1 (undiluted blood, 136 gametocytes/μl), 2/3 (91 gametocytes/μl) and 1/3 (45 gametocytes/μl). The isolate C with 784 gametocytes/μl in blood was treated to obtain the dilution factor 1/1 (undiluted blood, 784 gametocytes/μl), 1/3 (261 gametocytes/μl) and 1/8 (98 gametocytes/μl).

The reconstituted blood samples were provided in feeders for one hour to three to six days old females mosquitoes previously starved for 12hours. After exposure to blood meal, the unfed or partially fed females were removed and discarded, while the remaining fully engorged mosquitoes were kept in a bio secure room under standard conditions (27 ± 2°C, 70 ± 5% RH, 12:12 LD). The mosquitoes were given a 10% glucose solution on cotton wool after the blood meal.

### Mosquito midgut dissection

On the seventh day post blood meal (dpbm), 30 females exposed to each dilution factor of isolate A, about 24 (+/- 1) females exposed to each dilution factor of isolate B and about 10 (+/2) females exposed to each dilution factor of isolate C were dissected. Midguts were stained in a 1% mercurochrome solution and observed by microscopy to estimate the prevalence and intensity of oocysts in each group of exposed mosquitoes.

### Mosquito saliva collection and parasite DNA detection

A recently developed non-destructive sugar-feeding assay for parasite detection and estimating the extrinsic incubation period of *P. falciparum* in individual mosquito vectors was used [43]. Briefly, on the seventh dpbm, 20 to 40 females (median number = 30) exposed to each parasite isolate (A to C) and all experimental groups (dilution factors 1/1, 2/3, 1/3, 1/8) were individually placed in 28 ml plastic drosophila tubes with a cotton ball (15 mg/piece) soaked with 10% glucose solution placed on each tube gauze. Cotton balls were placed at 17:00 hrs on the tubes and removed the day after at 7:00 hrs. New cotton balls were placed daily on the mosquito tubes from 8 to 22 dpbm, then at 24 dpbm and finally every four days until the mosquito died. When removed, cotton balls were stored in sterile 1.5 ml Eppendorf tubes at −20 °C for further processing.

Upon death of all females used for saliva collection, DNA was extracted from head-thorax of each female using the DNeasy Blood and Tissue Kit system (Qiagen, Manchester, UK) according to the manufacturer’s instructions and parasite detection was carried out by qPCR, using *P. falciparum* mitochondrial DNA specific primers: qPCR-PfF 5’-TTA CAT CAG GAA TGT TTT GC-3’ and qPCR-PfR 5’-ATA TTG GGA TCT CCT GCA AAT-3’ [72]. For all females found positive by qPCR for *P. falciparum* in head-thorax extracts, genomic DNA from saliva in the cotton samples was also extracted using the same Qiagen protocol and the presence of sporozoites tested by the same qPCR protocol.

The DNA extracts from the heads-thoraces were tested 4 to 6 times each for the presence of parasite DNA by two different qPCR machines and the DNA extracts from cottons were run 3 times each by the same qPCR machine. Samples were considered positive for *P. falciparum* when at least one qPCR yielded a Ct < =38 and 75 <=Tm<= 80.

### Trait measurements

#### • Oocyst prevalence and intensity at 7 dpbm

For all experimental groups and for the three parasites isolates, 10 to 30 females were dissected for microscopic estimation of oocyst prevalence and intensity. Oocyst prevalence is the ratio of the number of mosquitoes with at least one oocyst out of all individuals dissected for each experimental group and each isolate. Oocyst intensity is the average number of oocysts in infected females for each experimental group and each parasite isolate.

#### • Sporozoite prevalence and intensity, extrinsic incubation period (EIP) and survival

The females placed in individual tubes to collect saliva in cotton balls were used to analyze the prevalence and intensity of sporozoites in carcasses (heads/thoraces) and in saliva, to measure the EIP of parasites and mosquito survival.

Sporozoite prevalence was expressed as the number of mosquito head/thoraces detected positive for *P. falciparum* by qPCR out of the total number of dissected head/thoraces for each treatment group and for each parasite isolate. Sporozoite intensity was expressed as the inverse of the mean number of threshold cycle during qPCR (the higher the 1/Ct value, the higher the infection intensity) for each treatment group and for each parasite isolate.

EIP was defined as the time between the infectious blood meal and the first day of positive molecular detection by qPCR of *P. falciparum* from the cotton wool where the female deposited saliva during sugar feeding.

Dead mosquitoes in the individual tubes in each experimental group were recorded every morning at 8:00 hrs for measurement of mosquito survival in each experimental group.

### Statistical analyses

All statistical analyses were performed in R (version 4.0.2). The effect of gametocyte density on oocyst and sporozoite prevalence was tested using logistic regression by generalized linear mixed models (GLMM, binomial errors, logit link; “lme4 package”), and its effect on oocyst and sporozoite density was tested using a negative binomial GLMM and a linear mixed model (“lme4” package), respectively. In these models, gametocyte density was set as both a fixed and a random slope effect and parasite isolate as a random intercept. EIP was analysed using two LMMs, the first specifying gametocyte density as a fixed and a random slope factor and parasite isolate as a random intercept, the second specifying sporozoite load in mosquito head/thorax as a fixed and a random slope factor and parasite isolate as a random intercept. We also investigated the effect of parasite isolate on EIP using a Cox’s proportional hazard regression model. The effect of infection status (infected vs uninfected) on mosquito survival was evaluated using a mixed Cox’s proportional hazard regression model (package “coxme”) with infection considered as a fixed effect and parasite isolate as a random intercept effect. Mosquito longevity was also analysed using two LMMs, the first specifying gametocyte density as a fixed and a random slope factor and parasite isolate as a random intercept, the second specifying sporozoite load in mosquito head/thorax as a fixed and a random slope factor and parasite isolate as a random intercept. Finally, we investigated the effect of parasite isolate on mosquito survival using a Cox’s proportional hazard regression model. For each model, the P < 0.05 statistical significance of the fixed effects was evaluated using the “Anova” function of the “car” package.

## Acknowledgments

We thank all volunteers for participating in this study as well as the local authorities for their support. We are very grateful to all the students and technicians at the IRSS/IRD who provided valuable assistance for the experiments of this study.

